# Oncogenic GLI1 and STAT1/3 activation drives immunosuppressive tryptophan/kynurenine metabolism via synergistic induction of IDO1 in skin cancer cells

**DOI:** 10.1101/2020.05.04.074757

**Authors:** Dominik P. Elmer, Victoria Strobl, Georg Stockmaier, Hieu-Hoa Dang, Markus Wiederstein, David Licha, Anna Strobl, Christina Sternberg, Suzana Tesanovic, Sandra Grund-Groeschke, Wolfgang Gruber, Florian Wolff, Richard Moriggl, Angela Risch, Roland Reischl, Christian G. Huber, Jutta Horejs-Hoeck, Fritz Aberger

**Affiliations:** Department of Biosciences, Cancer Cluster Salzburg, University of Salzburg, Salzburg, Austria; Department of Biochemistry, Christian-Albrechts-University Kiel, Kiel, Germany; Institute of Pathology, Unit of Laboratory Animal Pathology, University of Veterinary Medicine Vienna, Vienna, Austria; Institute of Animal Breeding and Genetics, University of Veterinary Medicine Vienna, Vienna, Austria

**Keywords:** Hedgehog signaling, GLI transcription factors, immune evasion, indoleamine 2,3-dioxygenase 1, Interleukin-6, Interferon-gamma, Signal Transducer and Activator of Transcription (STAT) proteins

## Abstract

Pharmacological targeting of Hedgehog (HH)/GLI has proven effective for certain blood, brain and skin cancers including basal cell carcinoma (BCC). However, limited response rates and the development of drug resistance call for improved anti-HH therapies that take into account synergistic crosstalk mechanisms and immune evasion strategies.

In previous work, we demonstrated that crosstalk of HH/GLI with pro-inflammatory Interleukin-6 (IL6) signaling drives BCC by promoting tumor cell proliferation [1]. In the present study, we screened for possible mechanisms of cancer immune evasion regulated by synergistic HH-IL6 signaling and identified the immunosuppressive enzyme indoleamine 2,3-dioxygenase 1 (IDO1) as a novel transcriptional target co-regulated by HH-IL6 signaling. Analysis of the *cis*-regulatory region of IDO1 by chromatin-immunoprecipitation revealed co-occupancy of this region by HH- IL6 induced GLI1 and STAT3 transcription factors along with active chromatin marks at the histone level. Elevated expression of IDO1 in human BCC with high-level HH and IL6 signatures supports the clinical relevance of our mechanistic data. Genetic inhibition of GLI1 expression prevented the induction of IDO1 expression in response to IL6/STAT3 and IFNγ/STAT1 signaling in human melanoma cells. Pharmacological targeting of HH signaling at the level of GLI proteins interfered with IDO1 expression and consequently prevented the production of the immunosuppressive metabolite kynurenine generated by active IDO1 from tryptophan. Further, inhibition of GLI1 enhanced the efficacy of the selective IDO1 inhibitor epacadostat. Of note, inhibition of HH/GLI signaling in melanoma cells not only reduced IDO1 expression but also interfered with the repression of T cell activation by attenuating IDO1/kynurenine-mediated immunosuppression. These data identify the immunosuppressive IDO1-kynurenine pathway as a novel pro-tumorigenic effector of oncogenic HH-IL6 and GLI-STAT cooperation. Our data suggest simultaneous pharmacological targeting of the HH/GLI, JAK/STAT and IDO1- kynurenine axis as rational combination therapy in skin cancers.

## Introduction

Uncontrolled activation of the Hedgehog (HH)/glioma-associated oncogene homolog (GLI) pathway can cause and promote the development and progression of several malignant diseases, of which basal cell carcinoma (BCC) with 3-4 million new cases per year in the US alone representing the most common cancer in the Western world [2]. During the past years, several small molecule inhibitors selectively targeting HH/GLI signaling were approved for the treatment of HH-driven cancers including advanced and metastatic basal cell carcinoma (BCC) and acute myeloid leukemia (reviewed in [3–8]). Despite these impressive advancements, frequent and rapid development of drug resistance, insufficient response and efficacy and severe adverse effects pose major challenges, calling for improved therapeutic strategies [9–12]. In light of the recent revolution in immuno-oncology, it is paramount also to identify possible immune modifying and suppressive mechanisms regulated by oncogenic HH/GLI signaling. This may allow for the development of rational combination therapies not only targeting the oncogenic HH/GLI signal but also interfering with putative immune evasion strategies to harness the potent anti-tumoral immune response.

HH/GLI signaling is a highly complex and intricately regulated pathway involving the release of repressive mechanisms, compartmentalization in the primary cilium, proteolytic processing and numerous post-translational modifications. In brief, HH/GLI signaling is initiated via binding of secreted HH protein to its receptor Patched (PTCH), which - via endocytotic removal of the HH- PTCH complex - allows the essential pathway effector Smoothened (SMO), a seven-pass transmembrane protein, to translocate into the primary cilium. There, active SMO initiates a signaling cascade that ultimately results in the formation of high-levels of active GLI transcription factors such as GLI1/2 driving HH target gene expression and tumor development (for detailed reviews see [7,13–17]).

In BCC, genetic loss of PTCH function or gain of function mutations in SMO results in ligand-independent irreversible HH/GLI signaling and skin carcinogenesis [9,18–20]. The strength and oncogenicity of HH/GLI signaling is further modified by synergistic cross-talk with other oncogenic pathways such as PI3K-AKT, MAPK, EGFR, DYRK or IL6/JAK/STAT [1,21–28]. A detailed understanding of critical cooperative interactions represents a crucial basis for the development of more efficacious combination therapies.

Our own group has recently shown that integration of HH/GLI and IL6/STAT3 signaling drives the expression of common HH-IL6 target genes and growth of BCC tumors [1]. Building on these data, we extended our analysis of HH-IL6 target genes with a focus on a putative role of HH-IL6 signaling in the suppression of the anti-tumoral immune response. Given the fact that HH/GLI-driven non-melanoma skin cancers display an exceptionally high mutational burden when compared to other cancer entities [29], we hypothesize that oncogenic HH/GLI signaling contributes to the establishment of an immunosuppressive microenvironment to prevent tumor elimination by the immune system. This concept was supported by the immune phenotype of murine BCC with immunosuppressive molecular and cellular signatures enriched in the tumor and its microenvironment (Grund-Gröschke, *et al*., 2019). In line, the potential therapeutic benefit of immunotherapy for BCC patients with immune checkpoint blockers (ICBs) has recently been reported in case studies, opening new opportunities for combination therapies targeting HH/GLI signaling and the immunosuppressive microenvironment [31–37].

In addition to the numerous immunosuppressive mechanisms involving immune checkpoint molecules, cytokines, hormones, growth factors and other smaller peptide signaling factors, several metabolites were identified as enhancers of cancer immune evasion. For instance, in cancer cells elevated expression of indoleamine 2,3-dioxygenase 1 (IDO1), the key enzyme responsible for extra-hepatic tryptophan catabolism, has been shown to cause local tryptophan starvation and to catalyze the production of the immunosuppressive metabolite kynurenine [38–40]. Both, tryptophan depletion and increased kynurenine levels, potently inhibit proliferation and are able to induce apoptosis of effector T cells and natural killer cells [41,42]. Of note, high levels of kynurenine generated by IDO1 establish a pronounced immunosuppressive milieu also via the induction and recruitment of immunosuppressive regulatory T cells (Tregs) and myeloid-derived suppressor cells (MDSCs) [43,44]. The potent immunosuppressive role of IDO1 in malignant settings has led to the development of several selective IDO1 inhibitors which are currently tested in clinical trials mainly as ICB adjuvant or pre-clinically in triple combination with radiotherapy [45–50].

Expression of IDO1 is intricately controlled by numerous inflammatory cytokines, often requiring a combination of two signals to reach physiologically relevant levels [39,44,51]. In melanoma, one of the strongest inducers of IDO1 expression is Interferon gamma (IFNγ) via activation of STAT1 [52,53]. Similarly, IL6 has been shown to induce IDO1 expression via JAK/STAT signaling in brain tissue, lung as well as ovarian carcinoma [54,55].

Based on our previous study on the synergistic cooperation of HH-IL6 signaling in non-melanoma skin cancer [1] and in light of the putative immunogenic nature of BCC [29], we screened for HH-IL6 target genes with a documented immunosuppressive function. We identified IDO1 as novel direct GLI1-STAT1/3 target gene cooperatively regulated by combined HH and cytokine signaling. Our functional studies using genetics, biochemical and immunological approaches support a model where combined GLI-STAT activation results in increased IDO1 levels and, as a consequence, in tryptophan starvation and enhanced kynurenine concentrations, leading to efficient inhibition of effector T cells. Thus, this study describes a novel oncogenic mechanism of HH/GLI signaling via IDO1-mediated tumor immune evasion with possible implications for combination therapies with IDO1 inhibitors and ICBs.

## Material and Methods

### Cell culture

All cells were maintained at 37 °C in an incubator with humidified atmosphere containing 5% CO_2_. Compounds and cytokines used for the treatment of cells are listed in supplementary Table S1.

In-house modified human keratinocyte (HaCaT) cell lines [56] with doxycycline (dox)-inducible GLI1 exogenous expression with and without Myc-tagged GLI1 had been generated before and were maintained and induced as described previously [26,57]. Human melanoma cell lines WM35 and WM793B were cultured in Dulbecco’s Modified Eagle’s Medium supplemented with 10% fetal bovine serum and 1% penicillin/streptomycin solution (Merck, Darmstadt, Germany). For generation of conditioned melanoma media for immune cell inhibition experiments tryptophan (Merck, Darmstadt, Germany) was added to the culture media to a final concentration of 200 µM to allow efficient kynurenine production in response to IDO1 expression.

All studies involving human immune cells were performed in agreement with the guidelines of the World Medical Association’s Declaration of Helsinki. In the case of anonymous blood cells discarded after plasmapheresis (buffy coats), national regulations do not request informed consent, thus no additional approval by the local ethics committee was required. Human peripheral blood mononuclear cells (PBMCs) were isolated from buffy coats of healthy, anonymous donors (provided by the Blood Bank Salzburg, Austria) and cultured as described previously [58]. Proliferation of CD4^+^ and CD8^+^ T cells was induced by stimulating cells with plate bound anti-CD3 [0.1 µg/mL] (Thermo Fisher Scientific, MA, USA) and soluble anti-CD28 [1 µg/mL] (BD Biosciences, NJ, USA).

### Microarray analysis

mRNA expression profiling has been described previously by [1]. Expression values were normalized to control and log2 transformed. The heatmap was generated using GraphPad Prism software (version 8.0.2, San Diego, CA, USA).

### RNA-seq data and clustering analysis

Fastq files of paired-end RNA-sequencing reads published by [9] were obtained from the Gene Expression Omnibus database (GEO accession: GSE58375). Reads were trimmed for adapter/primer sequences with Trim Galore (https://www.bioinformatics.babraham.ac.uk/projects/trim_galore/), a wrapper tool around FastQC (https://www.bioinformatics.babraham.ac.uk/projects/fastqc/) and cutadapt [59]. Subsequently, reads were aligned to the human reference genome GRCh37/hg19 with annotations from GENCODE (V30lift37) using the STAR software (version 2.7.0e, [60]) to obtain the abundance of reads per gene as counts. Count normalization to the library size and background noise elimination was conducted with the R package EdgeR (version 3.24.3, [61]) resulting in counts per million (cpm). Log2-transformed counts were subjected to data scaling and hierarchical clustering (Ward’s minimum variance method) based on signaling pathway gene signatures. Clustering results were depicted as heatmaps.

### Analysis of mRNA and protein expression

Total RNA was isolated following a standard phenol-chloroform extraction protocol using TRI-reagent (Merck, Darmstadt, Germany) with subsequent LiCl (Carl Roth, Karlsruhe, Germany) precipitation. cDNA was synthesized using M-MLV reverse transcriptase (Promega, Mannheim, Germany). Reverse transcription qPCR (RT-qPCR) runs were conducted as described previously [1]. The sequences of the RT-qPCR primers are provided in Supplementary Table S2. The synergy score was calculated as described by [62].

Standard protocols were used for SDS-PAGE and Western blot analysis of proteins. The antibodies used for protein detection can be found in Supplementary Table S3.

### Promoter analysis and chromatin immunoprecipitation

Putative GLI binding sites were predicted by the *D-Light* client-server software package using the matrix of consensus and non-consensus GLI binding site motives [63,64]. Information on STAT binding sites was retrieved from the ENCyclopedia Of DNA Elements (ENCODE) project [65] as described in [1].

ChIP was performed with the magnetic bead SimpleChIP Kit (Cell Signaling Technology, Boston, MA, USA) according to the manufacturer’s instructions (4 × 10^7^ cells per chromatin sample) followed by qPCR as described previously [1]. ChIP primer sequences and antibodies are listed in Supplementary Tables S2 and S3.

Quantitative methylation analysis of the CpG site closest to the STAT binding site (i) and GLI binding site (i) region was performed by bisulfite pyrosequencing as previously described in [1]. PCR and pyrosequencing primers are listed in Supplementary Table S2.

### RNA interference and lentiviral transduction

Production of lentiviral particles by 293FT cells using metafectene pro (Biontex Laboratories, Munich, Germany) and transduction of melanoma cells was performed following the protocol described in [66]. For RNA interference lentiviral particles were produced using following short hairpin RNA (shRNA) constructs purchased from the Mission TRC shRNA Library (Merck, Darmstadt, Germany): control shRNA (SHC002), shGLI1#1 (TRCN0000020486), shGLI1#2 (0000020488). After transduction cells were selected with puromycin (Merck, Darmstadt, Germany).

### Metabolite extraction and HPLC-MS analysis

For detailed method parameters see supplementary information. Briefly, metabolites were extracted from conditioned media by dilution (1:10) in ice-cold methanol (Merck, Darmstadt, Germany) containing 3-nitro-L-tyrosine [5 µM] (Merck, Darmstadt, Germany) as internal standard (ISTD) followed by centrifugation (10 min, 4°C, 13000 x g). Methanol extracts were diluted 1:5 in Milli-Q water (Merck, Darmstadt, Germany) and subjected to reversed phase HPLC-MS using an Accela 1250 HPLC system equipped with a Hypersil Gold aQ column and coupled to a QExactiveTM mass spectrometer (all from Thermo Fisher Scientific, MA, USA). Analyte separation was performed applying a multistep H2O - ACN gradient, metabolites were detected operating the QExactiveTM mass spectrometer in parallel reaction monitoring. Data were analyzed using the Thermo Xcalibur software (version 3.0.63, Thermo Fisher Scientific, MA, USA). Peak areas of tryptophan and kynurenine were normalized to the ISTD and used for relative quantification.

### Flow cytometry

PBMCs were stained with proliferation dye eFluor 450 [2 µM] (Thermo Fisher Scientific, MA, USA), seeded into anti-CD3 [0.1 µg/mL] (Thermo Fisher Scientific, MA, USA) pre-coated 48-well plates (2.5 × 10^5^ cells per well) and 50% conditioned melanoma or control medium were added (total volume 500 µL per well). PBMCs were then treated with anti-CD28 [1 µg/mL] (BD Biosciences, NJ, USA) and incubated for 72 h. After that, PBMCs were stained with the viability dye eFluor 780, anti-CD3-PE, anti-CD4-FITC and anti-CD8-BV510 (manufacturers and dilutions are listed in Supplementary Table S3), fixed in 4% paraformaldehyde (Merck, Darmstadt, Germany) and subjected to flow cytometric analysis on a BD FACS Canto II (BD Biosciences, NJ, USA). Flow cytometry data were analyzed with the FlowJo software (BD Biosciences, NJ, USA). The gating strategy is described in Supplementary Fig. S1.

### Statistical analysis

Statistically significant differences between two groups were tested using paired student’s t-tests, except for the analysis of human RNA-seq patient samples, where unpaired Welch’s t tests were used, with a confidence level of 95% for all analyses. Levels of significance were subdivided into following categories: \**P* < 0.0332, \*\**P* < 0.0021, \*\*\**P* < 0.0002, \*\*\*\**P* < 0.0001. Graphs and statistics were generated using GraphPad Prism software (version 8.0.2, San Diego, CA, USA). For each set of replicates the mean ± standard deviation (sd) indicated as error bars were depicted in the graphs.

## Results

### Synergistic HH/GLI and IL6/STAT3 signaling induces IDO1 expression in keratinocytes

In light of the exceptionally high mutational burden of BCC, we interrogated our HH-IL6 mRNA profiling data [1] for novel synergistically regulated HH-IL6 target genes with a known function in immunosuppression and cancer immune evasion. As displayed in the heat map in Figure 1A, we identified IDO1, a well-documented enzyme involved in cancer immune evasion, as a synergistically induced HH-IL6 target gene in human HaCaT keratinocytes. Addition of the JAK inhibitor panJAK inhibitor 1 efficiently suppressed the activation of IDO1 expression, suggesting JAK-dependent synergy of IL6 with HH/GLI signaling (Fig. 1A). We confirmed the gene expression profiling data by qPCR (Fig. 1B) and Western blotting (Fig. 1C). To induce GLI1 expression, cells were pre-treated with dox for 24h [50 ng/mL] and then treated with IL 6 [75 ng/mL] for another 24h. mRNA transcript levels of IDO1 showed a synergistic upregulation upon activation of combined HH and IL6 signaling with a synergy score of 0.47, indicative of more than additive activation of target gene expression [62] (Fig. 1B). Consistently, IDO1 protein levels were also strongly induced upon HH/GLI and IL6/STAT3 pathway activation (Fig. 1C).

**Fig. 1.**
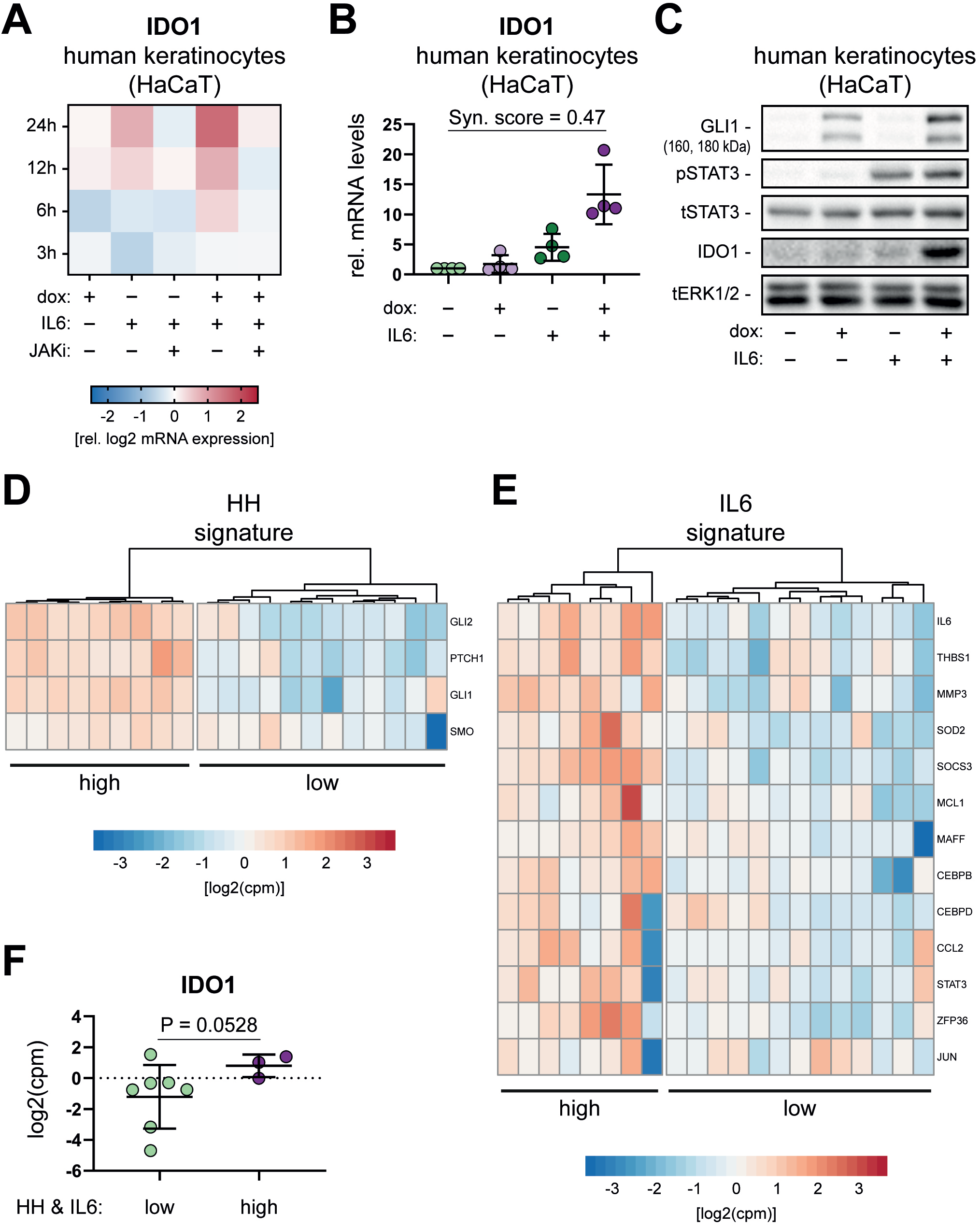
HH/GLI and IL6/STAT3 signaling cooperatively induce the expression of immunosuppressive IDO1. **(A)** Heatmap of log2 mRNA expression values of IDO1 measured by bead array expression profiling after 3, 6, 12 and 24 h in dox-inducible GLI1 human HaCaT keratinocytes in the presence or absence of IL6 [75 ng/mL] and panJAK inhibitor 1 [1 µM]. **(B)** IDO1 mRNA expression levels in HaCaT keratinocytes treated with dox to induce GLI1, with IL6 [75 ng/mL] or a combination of both relative to untreated controls as measured by qPCR. A synergy score of < 1.0 [62] reflects more than additive activation of IDO1 expression in response to combined doxGLI1/IL6 treatment. **(C)** Representative Western blot analysis of IDO1 expression in HaCaT keratinocytes treated for 48 h with dox to induce GLI1 expression, IL6 [75 ng/mL] or a combination of both for the last 18 h (*n* = 4). Total ERK1/2 (tERK1/2) protein expression was used as loading control. **(D-E)** Clustering analysis of log2(cpm) mRNA expression values of human BCC patient samples from the GEO database (GSE58375) using **(D)** HH/GLI and **(E)** IL6/STAT3 pathway signature genes (*n* = 21). **(F)** IDO1 log2(cpm) mRNA expression values of BCC patient samples grouped into HH-IL6 low (*n* = 7) or HH-IL6 high (*n* = 3) signaling activity according to the results of the clustering analysis shown in **(D)** and **(E)** (*n* = 21). Unpaired Welch’s t test was used for statistical analysis (\**P* < 0.0332). (cpm: counts per million; dox: doxycycline; JAKi: panJAK inhibitor 1; Syn. score: synergy score; p: phospho; t: total).

These *in vitro* data prompted us to investigate the cooperative up-regulation of IDO1 by the HH/GLI and IL6/STAT3 signaling pathways in an RNA-seq data set of human BCC patient samples published by [9] (GEO accession: GSE58375). Clustering analysis was performed in order to group the patient samples according to their HH/GLI and IL6/STAT3 pathway activity status (Fig. 1D, E). As shown in Figure 1F, BCC patients with highly active HH/GLI and IL6/STAT3 signaling showed elevated levels of IDO1 expression, although the data did not reach statistical significance. Taken together, these data support the notion that combined HH/GLI and IL6/STAT3 signaling synergistically activates the expression of the immunosuppressive enzyme IDO1 in human keratinocytes and human BCC.

### Mechanisms of *IDO1* promoter activation upon cooperation of HH/GLI and IL6/STAT3 signaling

Having shown that combined HH/GLI and IL6/STAT3 signaling cooperatively regulates IDO1 expression, we aimed to investigate the molecular basis of signal integration and IDO1 activation. We hypothesized that HH-IL6 signal integration converges at the level of the IDO1 *cis*-regulatory region. We, therefore, first screened for putative GLI binding sites within this region using the *in silico* binding site prediction tool *D-light* [63]. This resulted in the prediction of two putative GLI binding sites [64] (*GLI-bs(i)* and *GLI-bs(ii)*) at nucleotide position −2871 upstream of the transcription start site (TSS) and +1736 downstream of the TSS, respectively (Fig. 2A)). Binding of the transcription factor GLI1 was confirmed experimentally in human myc-tagged GLI1 inducible HaCaT keratinocytes by ChIP-qPCR. Myc-tagged GLI1 was significantly enriched on *GLI-bs(i)* and *GLI-bs(ii)* upon combined activation of HH and IL6 signaling (Fig. 2B). Furthermore, we retrieved data on two known *STAT-bs* in the IDO1 *cis*-regulatory region from the ENCODE database. As shown by a *D-light* analysis, both STAT binding sites are predicted as sequences recognized and bound by STAT3 as well as STAT1. Notably, *STAT-bs(i)* is located in close proximity to *GLI-bs(i)* (within 200bp; Fig. 2A). Using a ChIP-qPCR approach, we found STAT3 binding next to *GLI-bs(i)* in IL6-treated GLI1 expressing cells (Fig. 2C). Thus, we propose that combined binding and co-occupancy of the IDO1 *cis*-regulatory region by activating GLI and STAT transcription factors accounts for synergistic activation of IDO1 expression in response to HH-IL6 signaling.

**Fig. 2.**
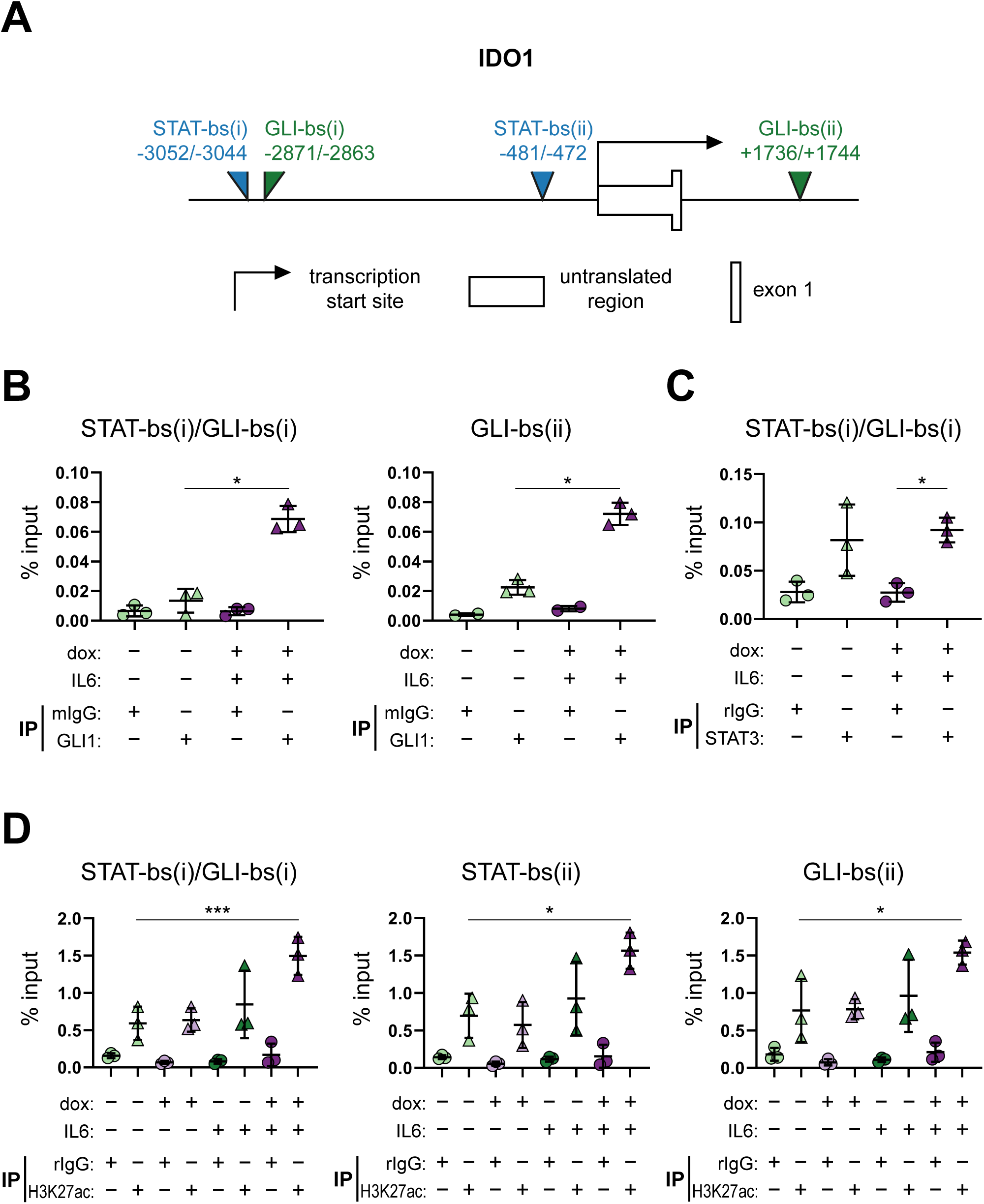
Combined activation of GLI1 and STAT3 results in co-occupancy of the IDO1 *cis*- regulatory region and promotes an active chromatin state. **(A)** Illustration of the IDO1 *cis*- regulatory region with *in silico* predicted GLI (green) and STAT (blue) binding sites with their position relative to the transcriptional start site (not drawn to scale). **(B-C)** Targeted ChIP analysis of GLI1 and STAT3 showing transcription factor binding to the following predicted sites in the IDO1 *cis*-regulatory region: **(B)** GLI1 binding to GLI-bs(i) as well as GLI-bs(ii) as evidenced by enrichment of the respective binding sequences upon GLI1 ChIP quantified by qPCR as percentage of input chromatin. **(C)** STAT3 binding to STAT-bs(i) immediately adjacent to GLI-bs(i). Human HaCaT keratinocytes expressing myc-tagged GLI1 under dox control were treated for 48 h with dox and 30 min with IL6 [75 ng/mL] to simultaneously activate STAT3 (*n* = 3). **(D)** ChIP analysis of active chromatin as determined by the level of H3K27 histone acetylation on STAT-bs(i)/GLI-bs(i), STAT-bs(ii) and GLI-bs(ii) expressed as percentage of input (*n* = 3). Cells were treated as described in (C). Student’s t test was used for statistical analysis (\**P* < 0.03; \*\*\**P* < 0.0002). (GLI-bs: GLI binding site; STAT-bs: STAT binding site, mIgG: mouse IgG; rIgG: rabbit IgG).

Additionally, we investigated whether the activity of HH/GLI alone or in combination with IL6/STAT3 signaling affects the epigenetic signature and landscape of the IDO1 *cis*-regulatory elements. To this end, we performed qPCR-ChIP experiments using antibodies specific for H3K27 acetylation, a marker for open active chromatin. Combined activation of both signaling pathways led to a significant enrichment of H3K27 acetylation, in contrast to the single treatments, at all three investigated binding regions: *STAT-bs(i)/GLI-bs(i), STAT-bs(ii)* and *GLI-bs(ii)* (Fig. 2D).

By contrast, combined HH/GLI-IL6 treatment did not alter the CpG methylation status in the region containing *STAT-bs(i)/GLI-bs(i)* (Fig. S2A, B).

### IL6/STAT3 as well as IFNγ/STAT1 mediated induction of IDO1 requires GLI1 in melanoma cells

To test and validate our findings in skin cancer models with a documented and pathophysiologically relevant role of IDO1 and oncogenic GLI expression, we switched our focus to human melanoma cells with JAK/STAT signaling dependent expression of IDO1 [27,43,46,52,67–73]. To address the possible role of endogenous, physiological oncogenic GLI activity, we performed RNA-interference (RNAi) mediated GLI1 inactivation in human BRAF^V600E^ mutated melanoma cells (WM35) and treated the cells with IL6 [75 ng/mL] for 18 h to activate STAT3 and IDO1 expression. *GLI1* knockdown and IDO1 levels were measured by qPCR analysis and/or Western blotting (Fig. 3A, B). In line with our data in human epidermal cells, *RNAi*-mediated inhibition of GLI1 expression with two different *siRNA* constructs hampered the induction of IDO1 expression in response to IL6 signaling on both the RNA and protein level (Fig. 3A, B).

**Fig. 3.**
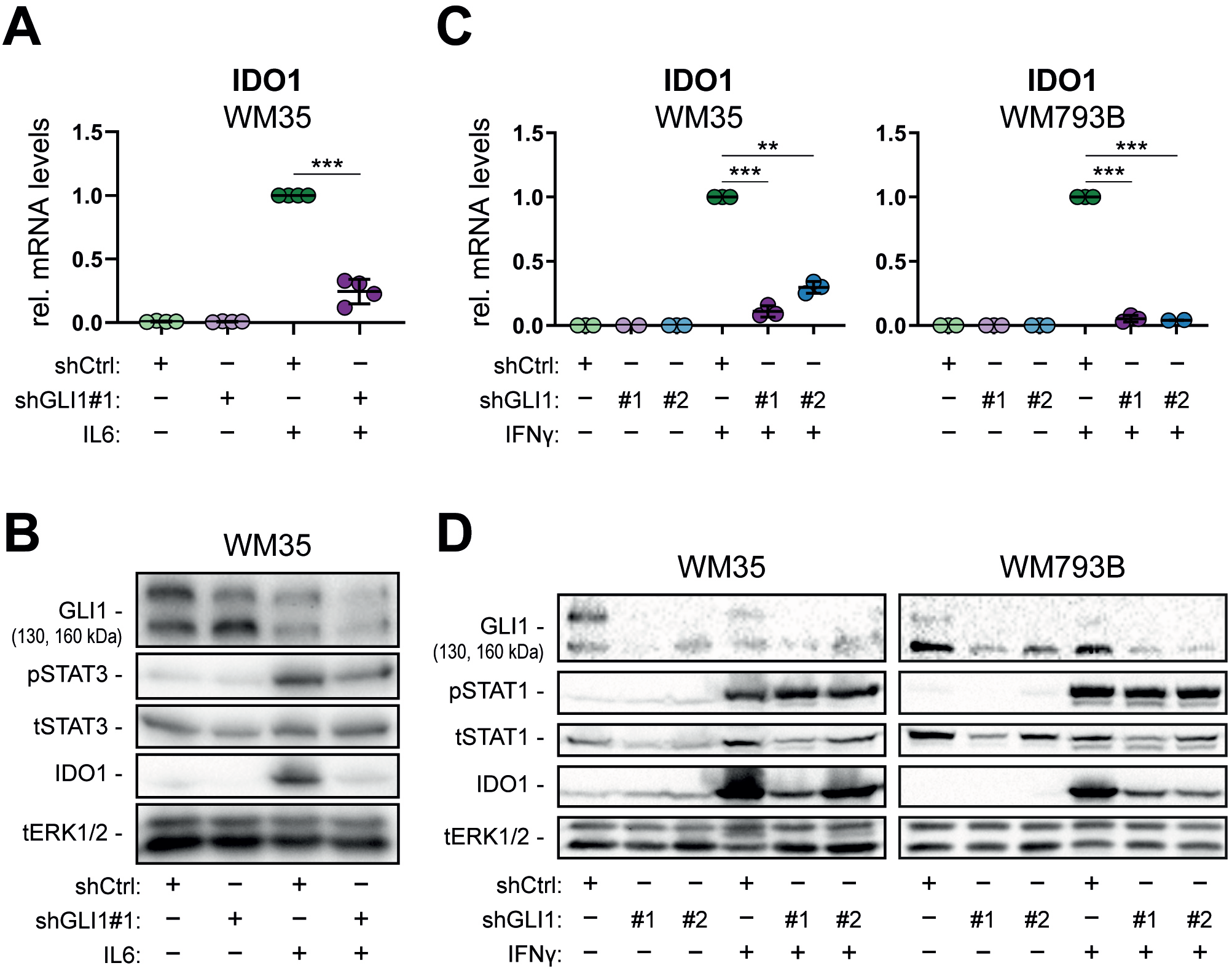
IL6 and IFNγ induced expression of IDO1 depends on GLI1 function in human melanoma cells. **(A)** qPCR analysis of IDO1 mRNA levels in WM35 melanoma cells treated with or without IL6 [75 ng/mL], and lentivirally transduced with shGLI1 (shGLI1#1) or control shRNA (shCtrl) (*n* = 3). (**B)** Western blot analysis of GLI1, total and phospho-STAT3 (tSTAT3 and pSTAT3) and IDO1 expression in WM35 melanoma cells treated as described in (A). **(C)** qPCR analysis of IDO1 mRNA levels in WM35 and WM793B melanoma cells treated with or without IFNγ (for 18 h with [10 ng/mL]), and lentivirally transduced with shGLI1 (#1 or #2) or control shRNA (shCtrl) (*n* = 3). **(D)** Western blot analysis of GLI1, total and phospho-STAT1 (tSTAT1 and pSTAT1) and IDO1 expression in WM35 and WM793B melanoma cells treated like in (C). Total ERK1/2 (tERK1/2) protein expression was used as loading control. Student’s t test was used for statistical analysis (\**P* < 0.035; \*\**P* < 0.002; \*\*\**P* < 0.0002; \*\*\*\**P* < 0.00001).

Since IFNγ/STAT1 signaling represents a well-known major and potent inducer of IDO1 expression [52,53], we also analyzed the requirement of GLI1 in settings where IDO1 expression is induced by the IFNγ/STAT1 axis. We performed *RNAi*-mediated knockdown of GLI1 in two *BRAF*^*V600E*^ mutant human melanoma cell lines (WM35 and WM793B) treated for 18 h with IFNγ [10 ng/mL]. Similar, yet more pronounced than in IL6-treated cells, genetic inhibition of GLI1 expression strongly reduced IFNγ-mediated induction of IDO1 expression in both melanoma cell lines (Fig. 3C). Of note, IDO1 protein levels strongly decreased by genetic targeting of GLI1 (Fig. 3D), supporting a crucial requirement of GLI1 in the activation of IDO1 expression in response to JAK/STAT signaling induced by IL6 or IFNγ.

### Pharmacological inhibition of the HH/GLI pathway reduces the capacity of melanoma cells to produce kynurenine

Following the genetic perturbation experiments, we next addressed whether pharmacological targeting of GLI is able to abolish the activation of immunosuppressive IDO1 expression. For this, we treated GLI1-expressing human melanoma cells (WM35) either with the FDA-approved Smoothened-inhibitor vismodegib [0.5 µM] or the GLI inhibitor Hedgehog Pathway Inhibitor 1 (HPI-1) [5 µM, 10 µM] for 6 h and then added IFNγ [10 ng/mL] for another 18 h to induce STAT1 activation and IDO1 expression. While vismodegib treatment neither reduced GLI1 protein levels (Fig. 4B) nor IDO1 expression (Fig. 4A, B), treatment with the GLI inhibitor HPI-1 profoundly reduced GLI1 protein (Fig. 4B) as well as IDO1 mRNA and protein expression (Fig. 4A, B), suggesting a crucial role of SMO-independent oncogenic GLI activity in the control of IDO1 expression.

**Fig. 4.**
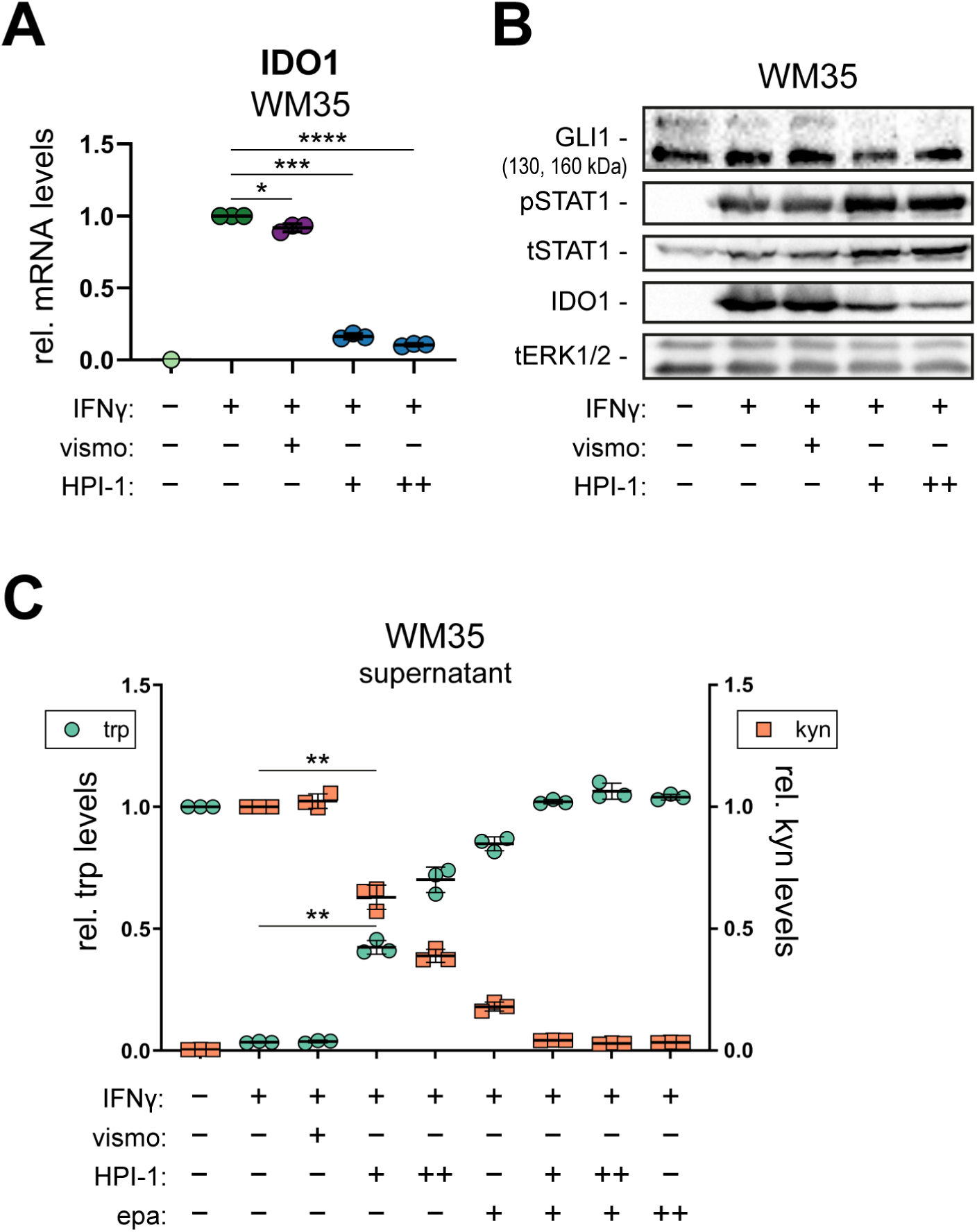
Pharmacological targeting of GLI but not of SMO prevents IFNγ-driven induction of IDO1 and production of immunosuppressive kynurenine metabolites. **(A)** qPCR analysis of IDO1 mRNA in WM35 melanoma cells treated with IFNγ [10 ng/mL], the GLI inhibitor HPI-1 or the SMO antagonist vismodegib (vismo), alone or in combination (n = 3). **(B)** Western blot analysis of GLI1, tSTAT1, pSTAT1 and IDO1 protein levels in WM35 melanoma cells treated for 24 h with solvent (control), vismo [0.5 µM], HPI-1 [5 µM] (+), [10 µM] (++) and/or IFNγ [10 ng/mL]. IFNγ was added following inhibitor pretreatment for 6 h. 130 kDa GLI1 represents a splice variant while the upper (∼160 kDa) band marks full-length GLI1, which we found to be more sensitive to HPI-1 than the 130 kDa form. Total ERK1/2 (tERK1/2) protein expression was used as loading control. **(C)** HPLC-MS analysis of tryptophan (green) and kynurenine (orange) levels in the supernatant of WM35 cells treated as indicated in the graph shown as fold change relative to solvent control or IFNγ, respectively (*n* = 3; vismo [0.5 µM]; HPI-1 [5 µM] (+), [10 µM] (++) epa [0.25 µM] (+), [1.5 µM] (++)). Student’s t-test was used for statistical analysis (\**P* < 0.035; \*\**P* < 0.002; \*\*\**P* < 0.0002; \*\*\*\**P* < 0.00001). (vismo: vismodegib; HPI-1: hedgehog pathway inhibitor 1; epa: epacadostat; trp: tryptophan; kyn: kynurenine).

To address whether changes of IDO1 protein levels in response to altered GLI/STAT activity affect tryptophan and kynurenine metabolite concentrations, we performed HPLC-MS measurements of the IDO1 substrate tryptophan and its product kynurenine in conditioned media of human melanoma cells (WM35). Correlating with IDO1 protein levels, kynurenine concentrations were highest in IFNγ/solvent or IFNγ/vismodegib treated cells, while tryptophan was quantitatively catabolized resulting in tryptophan depletion (Fig. 4C). By contrast, GLI inhibition by HPI-1 reduced kynurenine and rescued tryptophan levels in a concentration dependent manner. Low concentrations of the IDO1-specific inhibitor epacadostat [0.25 µM] partially inhibited kynurenine production, while complete inhibition of kynurenine production was achieved by the combination of low-dose epacadostat [0.25 µM] and HPI-1. Higher concentrations of epacadostat [1.5 µM] also blocked kynurenine production, indicating that the IFNγ-mediated production of kynurenine is mainly or even exclusively driven by IDO1 (Fig. 4C).

### Rescue of T cell proliferation upon pharmacological inhibition of the HH/GLI-IDO1 pathway in melanoma cells

To address the putative immunosuppressive role of the GLI/IDO1 axis on the activation of effector T cells known to play a key role in the anti-tumoral immune response, we investigated whether inhibition of GLI/IDO1 signaling is able to mitigate the immunosuppressive effects of these melanoma cells by reducing kynurenine levels. To that end, we generated conditioned media of melanoma cells treated with IFNγ (to induce IDO1 expression and kynurenine production) alone or in combination with inhibitors of GLI or IDO1. Conditioned media were then transferred to anti-CD3/anti-CD28 stimulated cells and T cell-proliferation was analyzed by flow cytometry. Representative flow cytometry data of CD4^+^ and CD8^+^ T cells treated with the different conditioned melanoma supernatants are shown in Fig. 5A. The results of three experiments with six individual donors are depicted in Fig. 5B, C. We found that melanoma cells treated with epacadostat alone did not cause a significant inhibition of CD4^+^ and CD8^+^ T cell proliferation, while HPI-1 slightly reduced T cell proliferation (Fig. 5B, C). By contrast, conditioned media from IFNγ-treated melanoma cells, which exhibit high levels of kynurenine due to strong IDO1 expression (see Fig. 4A-C), profoundly inhibited the proliferation of CD4^+^ as well as CD8^+^ T cells (Fig. 5B, C). Inhibition of IDO1 by epacadostat in IFNγ treated melanoma cells completely restored CD4^+^ and CD8^+^ T cell proliferation (Fig. 5B, C), demonstrating that T cell suppression by conditioned medium from IFNγ-treated melanoma cells is caused by increased IDO1 activity and high kynurenine levels (compare Fig. 4C; S3 A, B). Intriguingly and in line with our proposed model of GLI-dependent IDO1 expression, treatment of IFNγ- stimulated melanoma cells with the GLI inhibitor HPI-1 reinstated the proliferation of CD4^+^ and CD8^+^ T cells similar to epacadostat (Fig. 5B, C), providing functional evidence for a role of HH/GLI signaling as immunosuppressive driver via enhancing IDO1 expression and kynurenine production in the tumor microenvironment (summarized in Fig. 5D).

**Fig. 5.**
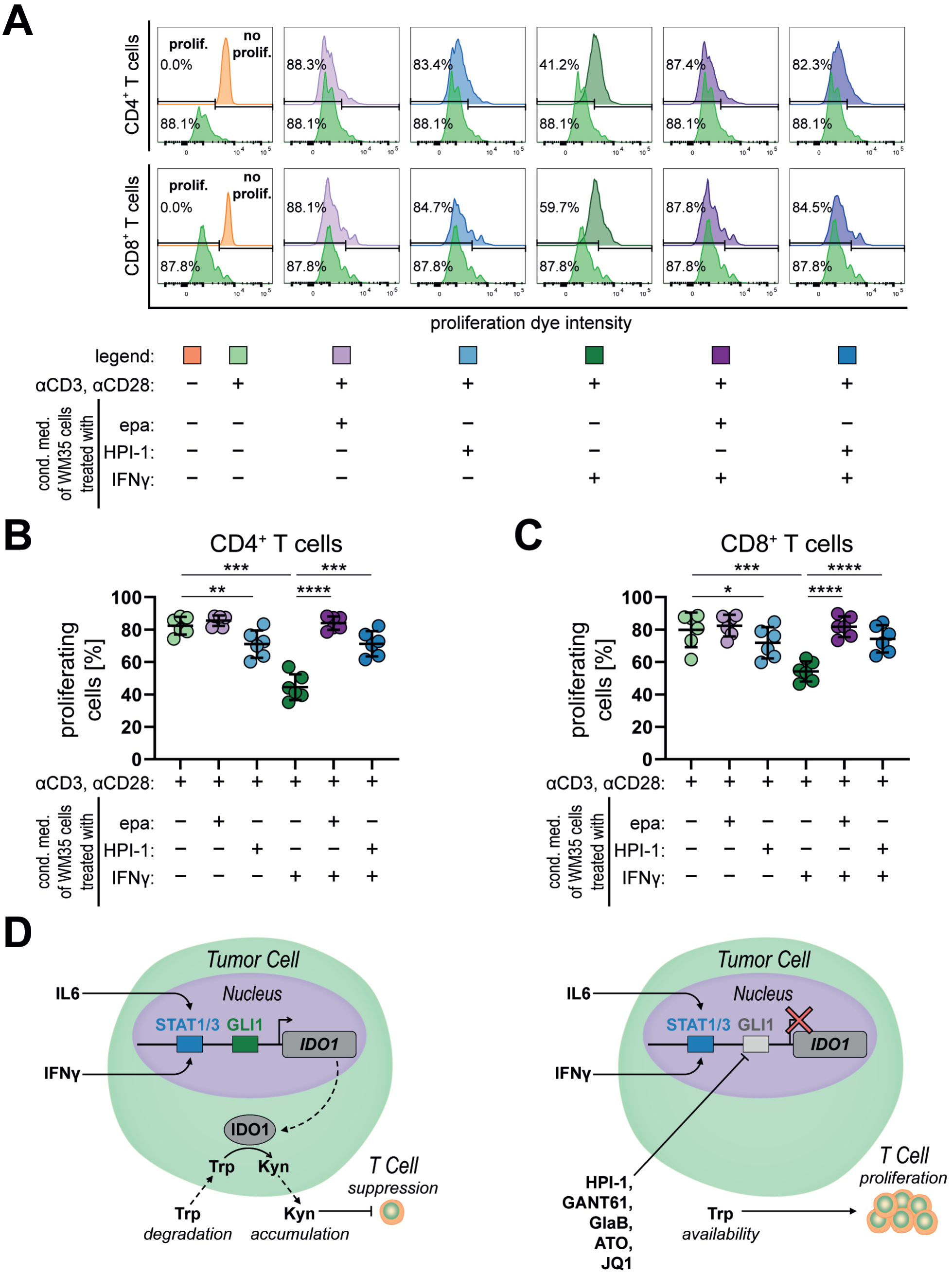
Pharmacological targeting of GLI reverts IDO1/kynurenine-dependent suppression of T cell proliferation. T cell proliferation in conditioned medium harvested from WM35 melanoma cells treated as indicated in the graph. **(A)** Representative flow cytometry analysis of anti-CD3/anti-CD28 induced proliferation of human primary CD4^+^ and CD8^+^ T cells in conditioned medium harvested from WM35 melanoma cells. **(B-C)** Flow cytometry analysis of the percentage of proliferating CD4^+^ (B) and CD8^+^ positive T cells (C) of anti-CD3/-CD28 stimulated PBMC cultures of different human donors each treated with a set of conditioned medium produced by treated WM35 cells (*n* = 6; epa [0.75 µM]; IFNγ [5 ng/mL]; HPI-1 [5 µM]). **(D)** Graphical summary of the proposed role of GLI-STAT signal integration in cancer immune modulation by IDO1 regulation. **Left:** Regulation of the IDO1 *cis*-regulatory region in the nucleus of tumor cells. GLI1 in cooperation with STAT1/3 mediates the transcriptional activation of IDO1 in the presence of inflammatory signals such as IL6 or IFNγ. Elevated IDO1 protein levels and enzyme activity lead to uptake and intracellular catabolism of tryptophan, which in turn generates high local levels of the immunosuppressive metabolite kynurenine in the tumor microenvironment. High levels of kynurenine efficiently suppress effector T cell activation, thereby interfering with the anti-tumoral immune response. **Right:** Pharmacological targeting of oncogenic GLI activator forms (e.g. by HPI-1, GANT61, GlaB, ATO or JQ1 [89–94]) abrogates the expression of IDO1 despite the presence of IDO1-inducing inflammatory signals such as IL6 or IFNγ. Therefore, GLI targeting results in reduced kynurenine and increased tryptophan concentrations in the tumor microenvironment, which helps reinstating T cell proliferation and anti-tumoral T cell activity. Student’s t-test was used for statistical analysis (\*\*\**P* < 0.0002; \*\*\*\**P* < 0.00001). (HPI-1: hedgehog pathway inhibitor 1; GANT61: GLI antagonist 61; GlaB: glabrescione B; ATO: arsenic trioxide; epa: epacadostat; trp: tryptophan; kyn: kynurenine).

To rule out that contaminating traces of inhibitors and/or inducers in the conditioned media account for changes in T cell proliferation independent of STAT/GLI-regulated IDO1 activity in melanoma cells, control supernatants containing the respective concentration of IFNγ, epacadostat and/or HPI-1 alone or in combination were tested on stimulated PBMCs (Fig. S3A, B). These control experiments clearly show that neither epacadostat [0.75 µM] nor IFNγ [5 ng/mL] affected CD4^+^ or CD8^+^ T cell proliferation (Fig. S3A, B), with HPI-1 showing only subtle, yet negative effects alone and in combination with epacadostat and IFNγ on T cell proliferation (Fig. S3A, B). As positive control demonstrating the repressive effect of the IDO1 product kynurenine, we show that treatment of PBMC cultures with kynurenine significantly inhibited CD4^+^ and CD8^+^ T cell proliferation (Fig. S3A, B), in line with its documented repressive effect on T cells [74–76].

## Discussion

In the present study, we report on a novel molecular mechanism of how oncogenic HH/GLI signaling can promote malignant growth by the activation of the tryptophan degrading immunosuppressive enzyme IDO1. Notably, we found that cooperative interactions of HH/GLI and pro-inflammatory IL6/STAT3 signaling drive high-level expression of IDO1. Activation of IDO1 expression represses effector T-cells via the production of the immunosuppressive metabolite kynurenine. On a molecular level, active GLI and STAT3 transcription factors bind to adjacent DNA-sequences in the *cis*-regulatory region, thereby synergistically inducing the transcription of IDO1, which is also accompanied by transcriptionally active histone marks in response to combined HH-IL6 signaling. Aside from a possible therapeutic relevance, activation of IDO1 in response to HH-IL6 signaling illustrates an example of how a well-documented pro-inflammatory and tumor-promoting pathway such as IL6 (Schaper and Rose-John, 2015; Taniguchi and Karin, 2014) is able to elicit in combination with HH/GLI a potent immunosuppressive signal. In combination, the tumor-promoting pro-inflammatory as well as the immunosuppressive activity may represent crucial mechanisms underlying the potent oncogenicity of both HH/GLI and IL6/STAT3 signaling.

In light of the promising preclinical and clinical data of trials with IDO1 inhibitors in combination with immune checkpoint inhibitors, our data on GLI/STAT-mediated regulation of IDO1 expression and immunosuppressive activity support the evaluation of novel treatments involving combinations of HH/GLI, JAK/STAT and IDO1 inhibitors for the efficient treatment of malignancies such as BCC. IDO1 inhibitors have been shown to reduce kynurenine concentrations and thereby enhance the efficacy of blocking immune checkpoints. However, recent phase III clinical trials failed to show a benefit of IDO inhibitors in combination with ICBs [79]. Whether this setback is due to a lack of predictive biomarkers and patient stratification remains to be addressed in future trials. Several possible reasons for this setback are discussed in the literature ranging from differences in the patient cohort to incomplete inhibition calling for a better understanding of the complex function and regulation of IDO1 in the tumor microenvironment [80–82].

The IDO1-kynurenine pathway has been shown to also shape the cellular landscape of the tumor immune microenvironment. For example, IDO inhibits crucial effector functions of cytotoxic T cells, as shown by reduced proliferation, cytokine release as well as degranulation and toxicity of CD8^+^ effector cells upon exposure to high IDO concentrations [83]. In addition, IDO-dependent tryptophan metabolites were shown to suppress proinflammatory Th1 responses [84], indicating that IDO attenuates important effector functions of CD4^+^ and CD8^+^ T cells. In line with these findings, we observe a clear inhibition of T cell proliferation upon culturing CD4^+^ and CD8^+^ cells with conditioned media derived from melanoma cells, which show high IDO expression. Kynurenine also enhances the formation of immunosuppressive Treg cells via binding to the aryl hydrocarbon receptor [85]. In this context, it is noteworthy that both murine and human BCC, which have been shown to express IDO1 [86], display increased Treg cell numbers [30]. Thus, it will be important to address in future studies the role of GLI/STAT and IDO1 activity in expansion and accumulation of Treg cells in the tumor immune microenvironment. Further, IDO1 activation in melanoma has been demonstrated to recruit myeloid derived suppressor cells (MDSCs) via enhancing Treg numbers, thereby contributing to immune evasion and resistance to immunotherapy [43]. Intriguingly, HH/GLI-induced BCC not only display increased numbers of Treg cells [30,87] but have also been shown to be infiltrated by immunosuppressive MDSCs [33]. Together, these data suggest that targeting of HH and JAK-STAT signaling in combination with IDO1 blockers and ICB may be a promising therapeutic approach for HH-associated malignancies, as such a combination would possibly not only target the oncogenic drivers but also re-establish the anti-tumoral immune response with more durable therapeutic effects.

IFNγ/STAT1 signaling is known to be both a crucial player in anti-tumoral immune responses as well as a strong activator of immunosuppressive IDO1 expression, particularly in melanoma [52,53,88]. We therefore investigated whether the HH/GLI signaling pathway also contributes to IFNγ/STAT1 regulated IDO1 expression. Similar to IL6/STAT3, genetic and pharmacological targeting of GLI1 showed that IFNγ/STAT1 requires functional GLI1 as a second signal for the full-blown induction of IDO1 in human melanoma cells. These findings add to our present understanding of the dual role of IFNγ in the anti-tumoral immune response and in cancer immune evasion and also exemplify how combinatorial signal integration events such as the cooperation of HH/GLI and IFNγ/STAT1 can affect the overall biological outcome.

In conclusion, our findings support the future evaluation of rational combination treatments involving HH/GLI and JAK/STAT inhibitors in combination with IDO1 blockers and/or ICBs to successfully target malignant growth and re-establish an efficient anti-tumoral immune response.

## Abbreviations

BCC: basal cell carcinoma
cpm: counts per million
dox: doxycycline
ENCODE: ENCyclopedia Of DNA Elements
GLI: glioma associated oncogene homolog
HH: Hedgehog
ICB: immune checkpoint blockade
ICBs: immune checkpoint blockers
IDO1: indoleamine 2,3-dioxygenase 1
IFNγ: Interferon-gamma
IL6: Interleukin-6
ISTD: internal standard
JAK: Janus Kinase
MDSCs: myeloid-derived suppressor cells
PBMCs: peripheral blood mononuclear cells
PTCH: Patched
RNAi: RNA interference
shRNA: short hairpin RNA
SMO: Smoothened
STAT: signal transducer and activator of transcription
Treg: regulatory T cell

## Acknowledgments

The authors are grateful to Drs Alexandra Kaser-Eichberger and Sandra Laner-Plamberger for dox-inducible MYC-tagged GLI1 cell lines, to Andrea Ramspacher for valuable input regarding figure design and discussions and to Mag. Christian Behensky, Sabine Siller and all other members of the Aberger group for continuous scientific and technical support. We acknowledge also the help of Martin Wipplinger and Helena Stadler with the RNA-interference experiments and methylation analyses.

## Authors’ contributions

DPE, VS, GS, H-HD, MW, DL, AS, CS and ST performed experiments. DPE, H-HD, MW, CS, FW, AR, JH-H, WG, RM, RR, CGH and FA designed experiments. All authors analyzed, discussed and interpreted data. DPE, SG-G and FA wrote the manuscript. All authors read and approved the final manuscript.

## Availability of data and materials

The authors declare that all data supporting the findings of this study are available within the article and its additional files and from the corresponding author upon reasonable request.

## Ethics approval

The use of anonymous buffy coats, which are blood cells that would be discarded after plasmapheresis (used for the isolation of PBMCs and provided by the Blood Bank Salzburg, Austria), does not require informed consent according to the national regulations in Austria. All human cells were handled according to the guidelines of the World Medical Association’s Declaration of Helsinki.

## Consent for publication

All authors have read the manuscript and agreed with publication of the study.

