## Supplementary Figures, Tables, Methods for "Oncogenic GLI1 and STAT1/3 activation drives immunosuppressive tryptophan/kynurenine metabolism via synergistic induction of IDO1 in skin cancer cells"

### Elmer et al. Supplementary Figures and Information

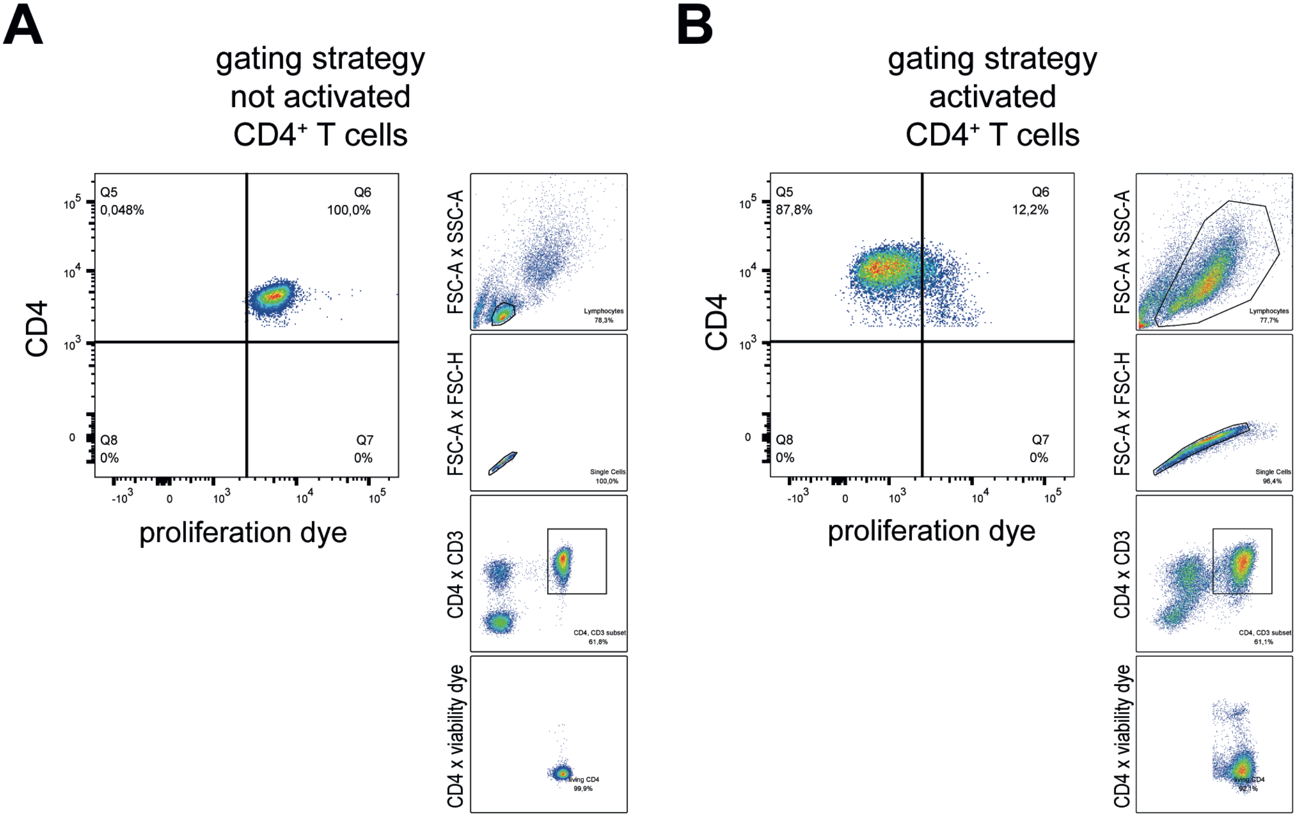


**Figure S1: Flow cytometry gating strategies.**

**(A)** Gating for the proliferation of CD4⁺ T cells in unstimulated PBMCs and **(B)** in αCD3, αCD28 activated PBMCs. **(A-B)** First the lymphocyte population was gated (FSC-A x SSC-A). Then doublet exclusion was performed (FSC-A x FSC-H). Next CD3 and CD4 double positive T cells were gated (CD4 x CD3). After that the living cells were gated from the CD3, CD4 double positive T cells (CD4 x viability dye). Final evaluation of the proliferation of the living CD3, CD4 double positive T cells (proliferation dye x CD4).

CD8⁺ T cells were gated accordingly.

### Elmer et al. Supplementary Figure 1

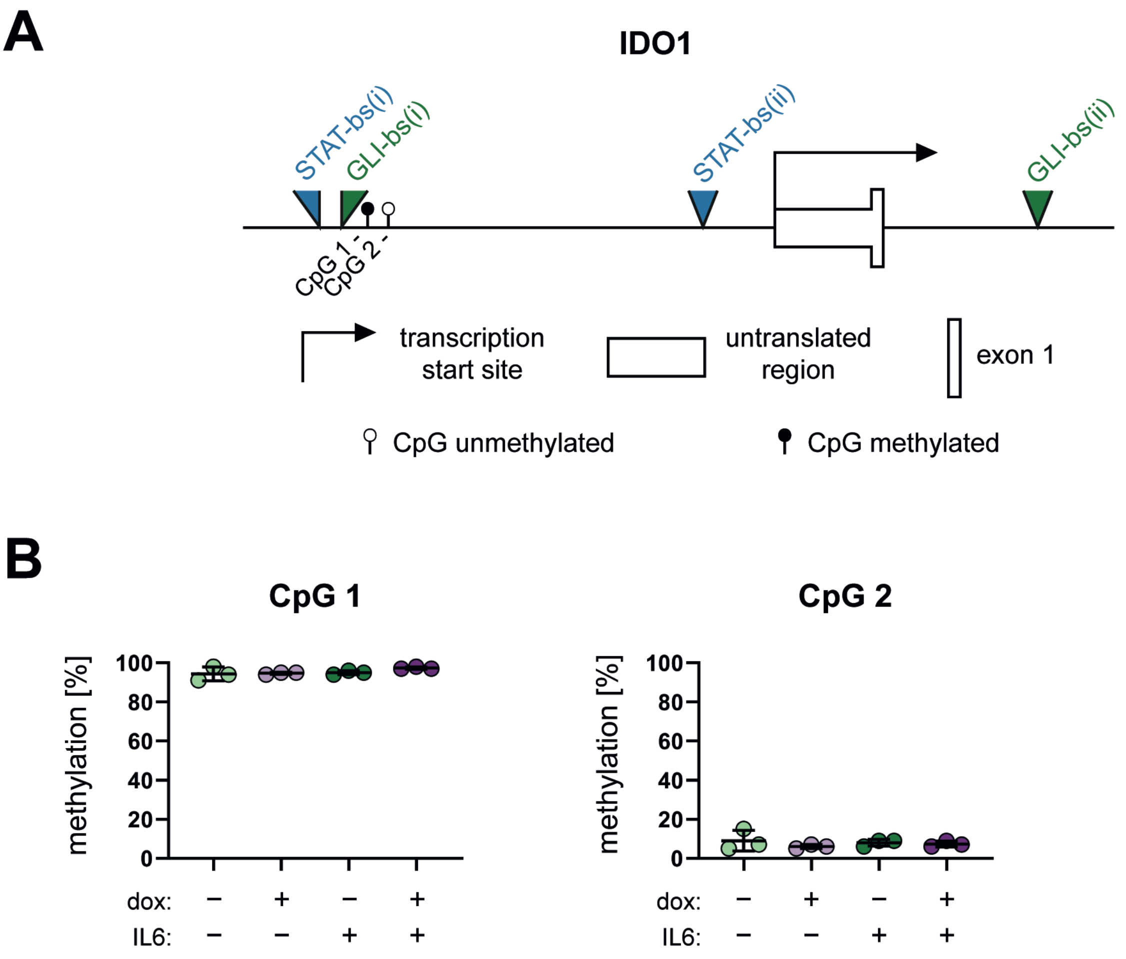


**Figure S2: CpG methylation analysis.**
**(A)** IDO1 *cis*-regulatory region with CpG 1 and CpG 2 next to GLI-bs(i) indicated.
**(B)** Bisulfite pyrosequencing analysis of the methylation status of CpG 1 and CpG 2 in dox-inducible human keratinocytes (HaCaT).

Elmer et al. Supplementary Figure 2

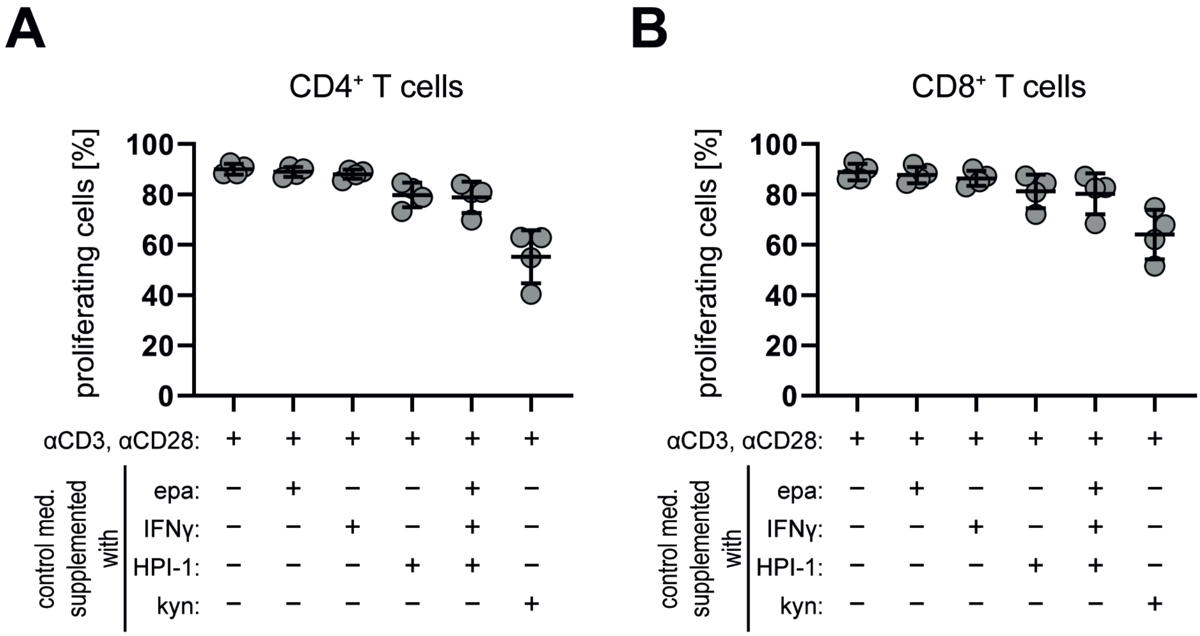


**Figure S3: PBMC treatment with control media.**

PBMCs were activated and treated with control media for 72h. The final treatment concentrations were: epa [0.75 μM], IFNγ [5 ng/mL], HPI-1 [5 μM], kyn [100 μM]. Flow cytometric analysis of the proliferation of $CD4^+$ **(A)** and $CD8^+$ T cells **(B)**.

Elmer et al. Supplementary Figure 3

Elmer et al. Supporting Information – Tables (S1 - S3):

**Table S1: Compounds and cytokines**

| **Substance** | **Manufacturer** | **Cat.No.** |
|---|---|---|
| Dimethyl sulfoxide (DMSO) | Merck | D2650 |
| Vismodegib (vismo) | LC Laboratories | V-4050 |
| Hedgehog pathway inhibitor 1 (HPI-1) | Merck | H1542 |
| Epacadostat (epa) | Selleckchem | S7910 |
| Doxycycline (dox) | Merck | D9891 |
| Interleukin 6 (IL6) | Peprotech | 200-06 |
| Interferon gamma (IFNγ) | Immuno Tools | 11343534 |
| L-tryptophan (trp) | Merck | T8941 |
| L-kynurenine (kyn) | Merck | K8625 |

**Table S2: Nucleotide sequences of primers**

| Target | Fw/Rv | Sequence (5'-3') | Company | Application |
|---|---|---|---|---|
| RPLP0 | Fw | GGCACCATTGAAATCCTGAGTGATGTG | Microsynth | RT-qPCR |
| | Rv | TTGCGGACACCCTCCAGGAAGC | | |
| GLI1 endogenous | Fw | TCTGGACATACCCCACCTCCCTCTG | Microsynth | RT-qPCR |
| | Rv | ACTGCAGCTCCCCCAATTTTTCTGG | | |
| GLI1 | Fw | GCCGTGCTAAAGCTCCAGTGAACACA | Microsynth | RT-qPCR |
| | Rv | TCCCACTTTGAGAGGCCCATAGCAAG | | |
| IDO1 | Fw | TGGGAAGACCCAAAGGAGTTTGCAG | Microsynth | RT-qPCR |
| | Rv | GACCAGAGCTTTCACACAGGCGTCA | | |
| IDO1 STAT-bs(i)/GLI-bs(i) | Fw | GATCCTTTGCCATTCAGCTACTAA | Microsynth | ChIP-qPCR |
| | Rv | CCCAGTAATTCCCCTGTTTGA | | |
| IDO1 STAT-bs(ii) | Fw | CGTTGTTCCATGTCTTCACTT | Microsynth | ChIP-qPCR |
| | Rv | GAGGCTAATACTTTGGGAATC | | |
| IDO1 GLI-bs(ii) | Fw | GGCCAGTATGAGCCTAAGCAG | Microsynth | ChIP-qPCR |
| | Rv | TTAAGAGAGGCAGTGTGGAATAATG | | |
| IDO1 STAT-bs(i)/GLI-bs(i) region | Fw | GGAATTATTGGGGTTTAGAAAAGGA | Merck | PCR (bisulfite converted DNA) |
| | Rv | [Btn]-ACAAACTTTAATCATTCTTCCTATAACAT | | |
| | - | GTTGGTTATTTTAAATTAAAGGT | Merck | Pyrosequencing |

Fw: forward; Rv: reverse; [Btn]: Biotin tag

**Table S3: Antibodies (Western blot, ChIP, Flow cytometry)**

| Antibody | Dilution | Manufacturer | Cat.No. | Application |
|---|---|---|---|---|
| GLI1 | 1:1000 | Cell Signaling Technology | 2534 | Western blot |
| STAT3 | 1:6000 | BD Biosciences | 610189 | Western blot |
| Phospho-STAT3 | 1:1000 | Cell Signaling Technology | 9131 | Western blot |
| STAT1 | 1:2000 | BD Biosciences | 610115 | Western blot |
| Phospho-STAT1 | 1:1000 | Cell Signaling Technology | 9167 | Western blot |
| IDO1 | 1:1000 | Cell Signaling Technology | 86630 | Western blot |
| ERK1/2 | 1:1000 | Cell Signaling Technology | 9102 | Western blot |
| Anti-rabbit, HRP | 1:3000 | Cell Signaling Technology | 7074 | Western blot |
| Anti-mouse, HRP | 1:3000 | Cell Signaling Technology | 7076 | Western blot |
| myc-tag | 1:100 | Cell Signaling Technology | 2276 | ChIP |
| Normal mouse IgG | same as test ab | Cell Signaling Technology | 5415 | ChIP |
| STAT3 | 5 µg/10 µg chromatin | Santa Cruz Biotechnology | sc-482X | ChIP |
| Normal rabbit IgG | same as test ab | Santa Cruz Biotechnology | sc-2027X | ChIP |
| H3K27ac | 1:100 | Cell Signaling Technology | 8173 | ChIP |
| Normal rabbit IgG | same as test ab | Cell Signaling Technology | 2729 | ChIP |
| eFluor 450 | 1:5000 | Thermo Fisher Scientific | 65-0842-85 | Flow cytometry |
| eFluor 780 | 1:3000 | Thermo Fisher Scientific | 65-0865-14 | Flow cytometry |
| CD3 (PE) | 1:25 | Immuno Tools | 21850034 | Flow cytometry |
| CD4 (FITC) | 1:50 | Thermo Fisher Scientific | 11-0048-42 | Flow cytometry |
| CD8 (BV510) | 1:50 | BD Biosciences | 563919 | Flow cytometry |

**Supplementary experimental details:w**

#### HPLC-MS Method for the targeted analysis of kynurenine and tryptophan

##### Sample preparation

In order to pellet proteins and to extract metabolites, 900 µL ice-cold methanol containing 5.0 µM 3-nitro-L-tyrosine as an internal standard (Sigma-Aldrich, Austria) were added to 100 µL of stored supernatant cell media. Proteins were pelleted by centrifugation at 18620 g and 4.0 °C for 10 minutes. The supernatant was used for HPLC-MS analysis after dilution with ultrapure water at a ratio of 1:5.

##### Targeted HPLC-MS analysis

An ultra-high performance liquid chromatography (UHPLC) system consisting of an Accela 1250 pump, a Column Oven 300 (all from Thermo Fisher Scientific, Bremen, Germany), and an LC PAL DLW Option Autosampler with a 100 µL syringe (from CTC Analytics AG, Zwingen, Switzerland) was coupled to a hybrid quadrupole-Orbitrap mass spectrometer (Model Q Exactive$^{TM}$; Thermo Fisher Scientific) equipped with a heated electrospray ion source operating in the positive ion mode. Prior to injection into the HPLC-MS system the injection order was randomized.

In order to monitor potential carry over, a blank run with Millipore water as an injection solute was performed after every third injection of sample. This summed up to 31 sample runs and 12 blank runs.

##### HPLC settings

For RP-HPLC separations a 100 x 2.1mm inner diameter Hypersil Gold aQ column (Thermo Fisher Scientific) packed with 1.9 µm octadecyl silica particles was applied. For column protection, a 4.0 x 3.0 mm inner-diameter $C_{18}$ Security Guard pre-column (Phenomenex, Torrance, California, USA) was installed.

Mobile phase A and B were Millipore water and acetonitrile, respectively, both containing 0.10 % formic acid (Sigma-Aldrich). The stepped gradient HPLC method started with holding 4.0 % B for 0.50 minutes, followed by the first gradient to 7.0 % B in 2.0 minutes and a second one to 50% B in 1.5 minutes. After washing for 1.0 minutes at 100 % B, the column was re-equilibrated to starting conditions for 3.0 minutes, resulting in a total run time of 8.0 minutes.

A flow rate of 0.30 µL min$^{-1}$ and an injection volume of 2.70 µL were applied. The column temperature was held constant at 30 °C.

##### MS-settings

A Parallel Reaction Monitoring (PRM) method was applied, with the orbitrap resolution set to 17500, an AGC target of 2e5 and a maximum injection time of 100 ms. The isolation window was set to 1.0 m/z and signals were acquired in profile mode. A scheduled inclusion list with distinctive normalized collision energies (NCEs) was applied for $[M+H^{+}]$ of kynurenine (209.0920 m/z), tryptophan (205.0970 m/z) and the internal standard 3-nitro-L-tyrosine (227.0660 m/z) as depicted in table 1.

As tune parameters a voltage of 3.2 kV, a capillary temperature of 350°C and an S-lens level of 55 were chosen. Sheath gas and auxiliary gas flow rates were set to 45 and 10 arbitrary units, respectively.

**Scheduled inclusion list, showing applied time windows and normalized collision energies for kynurenine, tryptophan and nitrotyrosine**

| Mass [m/z] | Polarity | Start [min] | End [min] | NCE |
|---|---|---|---|---|
| 209.0920 | Positive | 0.00 | 2.00 | 35 |
| 227.0660 | Positive | 0.00 | 2.00 | 35 |
| 205.0970 | Positive | 0.00 | 2.00 | 28 |
| 209.0920 | Positive | 2.00 | 2.65 | 35 |
| 227.0660 | Positive | 2.50 | 3.50 | 35 |
| 205.0970 | Positive | 3.50 | 5.00 | 28 |
| 209.0920 | Positive | 5.00 | 8.00 | 35 |
| 227.0660 | Positive | 5.00 | 8.00 | 35 |
| 205.0970 | Positive | 5.00 | 8.00 | 28 |

**Data evaluation**

Acquired .raw files were evaluated via the QuanBrowser in Thermo Xcalibur 3.0.63. The seven most intense fragment ions were used for quantification of tryptophan (118.0653, 132.0815, 144.0803, 146.0604, 159.0908, 170.0593, 188.0712 m/z) and kynurenine (94.0660, 120.0445, 136.0764, 146.0598, 150.0555, 174.0542, 192.0661 m/z), while only four fragment ions were consulted for the quantification of the internal standard 3-nitro-L-tyrosine (133.0522, 164.0337, 168.0289, 181.0606 m/z). For the quantification of all three analytes, a mass tolerance of 10 ppm, 5 smoothing points and a baseline window of 100 were applied. For tryptophan and kynurenine area noise factor and peak noise factor were set to 1, while for 3-nitro-L-tyrosine values of 2 and 10 were used, respectively. A signal to noise ratio of 9:1 was considered as limit of quantification. Calculated peak areas of tryptophan and kynurenine were normalized to the peak area of 3-nitro-L-tyrosine for each run.
